# Metabolic responses to cold: thermal physiology of native common waxbills (*Estrilda astrild*)

**DOI:** 10.1101/2024.01.18.576192

**Authors:** Cesare Pacioni, Marina Sentís, Anvar Kerimov, Andrey Bushuev, Colleen T. Downs, Luc Lens, Diederik Strubbe

**Author notes:** Corresponding author ORCID: 0000-0001-7773-9151.

## Abstract

Ecophysiological studies of invasive species tend to focus on captive individuals and their invasive range. However, the importance of gaining a thorough understanding of their physiology in their native range, where autecological knowledge is limited, has played a crucial role in assessing species ecophysiological responses to the often novel environmental conditions encountered in their invasive range. Here, we investigated the ecophysiological characteristics of a population of the wild-caught common waxbill (*Estrilda astrild*), a successful global invader, in part of its native range (South Africa). We investigated how this species adjusts its resting metabolic rate over a range of temperatures to identify its thermoneutral zone (TNZ) as an indicator of a species’ thermal tolerance. The observed TNZ curve predominantly followed the classic Scholander-Irving model, with metabolic rates increasing linearly at temperatures outside the TNZ. However, we found an inflection point at moderately cold temperatures (16°C) where the common waxbill began to decrease its metabolic rate. This finding highlights the potential use of an energy-saving strategy as an adaptive response to cold, such as facultative hypothermic responses through a reduction in body temperature, which may explain their success as an invasive species. We argue that although metabolic infection points have been repeatedly identified in studies of TNZ, the specific mechanisms behind metabolic down-regulation at low temperatures remain underexplored in the literature. We therefore suggest that future research should focus on investigating body temperature variation, with particular emphasis on its potential contribution to metabolic adaptation in colder environments.

## Introduction

Global climate change, characterized by rising temperatures and more frequent extreme weather events, is putting up to one out of six species at risk of extinction as they face unprecedented challenges in responding to rapid environmental change (Urban, 2015). Furthermore, the introduction of alien species poses a significant additional threat to global biodiversity, ecosystem services and economic resources (Bongaarts, 2019; Castro-Díez et al., 2019; Diagne et al., 2021; Pyšek et al., 2020; Shirley & Kark, 2009). According to the latest IPBES Invasive Alien Species Assessment (2023), at least 37.000 established alien species have been introduced by human activities, of which 3.500 are also invasive. Changes in global temperature and precipitation regimes are likely to amplify the impacts of invasive species (Dukes & Mooney, 1999; Walther et al., 2009), for example, by allowing populations of introduced species that are not presently invasive to become invasive (i.e., spread and cause impacts; Hellmann et al., 2008; Mainka & Howard, 2010). For both the effective conservation of native species and the management of invasive alien species, it is crucial to accurately assess how species will respond to new environmental conditions. In this context, knowledge of the physiological mechanisms used by species to adapt to changing climatic conditions can greatly improve ecological predictions of species range dynamics (Huey et al., 2021).

Understanding how animals thermoregulate to buffer unfavorable thermal fluctuations in their environment, and what their thermoregulatory limits are, have long been central questions in animal ecophysiology (Bozinovic et al., 2011). Thermoregulation in endotherms is generally non-linear and is thought to follow the classic Scholander-Irving model of endothermic homeothermy (Scholander et al., 1950), which includes a thermoneutral zone (TNZ). The latter defines the range of ambient temperatures within which an endotherm can maintain its body temperature without increasing its metabolic rate above the basal metabolic rate (BMR) required to maintain basic life-sustaining functions (McNab, 2012). The TNZ of a species is characterized by two critical temperatures: the lower critical temperature (LCT) and the upper critical temperature (UCT). Both mark the point at which the metabolic rate begins to rise above the BMR to maintain normothermia. Several studies have used the concept of TNZ as an indicator of a species’ long-term ability to tolerate thermal variation. For example, Khaliq et al. (2014) showed that around 15% of bird species presently experience maximum ambient temperatures above their UCT, and this rises to over 35% under climate change scenarios. This suggests that birds will increasingly face thermoregulatory constraints on fitness-related traits, such as activity levels, reproduction, and survival. For example, Milne et al. (2015) studied 12 bird species in South Africa and identified potential links between climate warming and population declines in several passerine species associated with fynbos habitat, such as the Cape rockjumper (*Chaetops frenatus*), and attributed these declines to the birds’ limited tolerance to higher temperatures.

A better understanding of thermoregulatory strategies and capacities can also contribute to more accurate risk assessment of invasive species, as they often expand their niche into novel climates (Liu et al., 2022). For example, ring-necked parakeets *(Psittacula krameri*) have successfully colonized many European cities, which are considerably colder than their native range (Strubbe et al., 2015). The same applies to several members of the family Estrildidae, which is notable for establishing numerous invasive bird populations (Stiels et al., 2015). This group of small, finch-like birds comprises 138 species (Winkler et al., 2020) and occupies diverse habitats across Africa, southern Asia, and Australasia, with the highest species concentrations occurring in the tropics (Goodwin, 1982). While several estrildid finches are classified as regional agricultural ‘pests’ (Gleditsch & Brooks, 2020), they are widely traded and have been described as the ‘single most important avicultural family’ (Ribeiro et al., 2020). With 81 introduced species worldwide (Dyer et al., 2017), it is considered the most successful non-native family of birds among tropical passerines (Lever, 2005), with most of the species traded as pet birds (Cardoso & Reino, 2018; Reino et al., 2017).

Due to their widespread popularity as pet cage birds (Reino et al., 2017), most studies on estrildid physiology have focused on captive individuals, while relatively few have examined the ecophysiology of free-ranging individuals (Allen & Hume, 2001; Cooper et al., 2019, 2020; Gerson et al., 2019; Sheldon & Griffith, 2018). Pacioni et al. (2023) and Sentís et al. (2023) studied the energetic metabolism of captive common waxbills (*Estrilda astrild*, 7-9 g), but no further information is available on the free-ranging individuals. Two closely related Aftrotropical estrildid finches, which also have established non-native populations (Ascensão et al., 2021), have also only been studied in captivity. Marschall and Prinzinger (1991) studied the thermal physiology of five estrildid species, including the orange-cheeked waxbill (*E. melpoda*), and showed that each of these birds had different physiological adaptations to their specific habitats in different tropical climates. Cade et al. (1965) and Lasiewski et al. (1964) studied the physiology of captive black-rumped waxbills (*E. troglodytes*), but found contrasting results regarding the species’ TNZ. Lasiewski et al. (1964) also claimed that when properly fasted, waxbill metabolic rates measured during the day (in darkened metabolic chambers) and during the night were similar. Stephens et al. (2001) came to the same conclusion for captive orange-cheeked waxbills. These findings contrast with review studies (e.g., Aschoff & Pohl, 1970) which suggest that daytime avian metabolic rates are on average 20-25% higher than nighttime rates, highlighting the need for further investigation of diurnal variation in metabolic rates in estrildid species.

Studying invasive species solely in captivity or in their invaded range might overlook crucial information about their ability to adjust to specific environmental conditions (Boardman et al., 2022). For example, it has been shown that a broader understanding of species in their native distribution range is essential for assessing how species may respond to the often novel environmental conditions they encounter in their introduced distribution range (Stuart et al., 2023). However, in contrast to the increasing knowledge of invasive species in their non-native range, autecological knowledge of many invasive species in their native range is still minimal (Ros et al., 2016). The same applies for the common waxbill, for which the distribution and dispersal of well-studied invasive populations in Iberia have been found to be strongly influenced by climate and habitat gradients (Sullivan & Franco, 2018), but accurate prediction of invasion risk is difficult (Stiels et al., 2015). In an attempt to improve accuracy, Strubbe et al. (2023) used thermal physiological approaches to model the species’ potential European distribution range expansion, but faced challenges in inferring key functional traits because of a lack of empirical data from wild individuals, forcing them to rely on allometric predictions. Moreover, while experiments on captive birds have value, the non-natural biotic and abiotic conditions to which animals are exposed can lead to important ecophysiological changes (Beaulieu, 2016). Prioritizing research on wild animals, as well as on their native distribution range, is therefore crucial for gaining ecologically relevant insights into species’ potential for distributional expansion. Therefore, here we investigated the ecophysiological characteristics of wild-caught common waxbills in a part of their native range (South Africa). Specifically, we assessed how this species adjusts its ρ-(resting) phase metabolic rate over a range of temperatures to identify its thermoneutral zone. We also investigated α-(active) phase metabolic rate patterns, predicting that (i) fasted birds would have a lower resting metabolic rate than non-fasted birds, and (ii) the α-phase metabolic rates of fasted birds would be higher than the ρ-phase metabolic rates. By filling this gap in the understanding of how wild-caught common waxbills in a part of their native distribution range physiologically adjust to temperature variation, our research will provide crucial data that could potentially be incorporated into models aimed at predicting invasion dynamics and associated risks.

## Methods

### Study area and bird capture and maintenance

The study was conducted in South Africa (Pietermaritzburg, KwaZulu-Natal) near the Darvill Wastewater Treatment Site (N −29.60, E 30.43). Between September 30 and November 4 (2022), free-ranging common waxbills were attracted by song playback and captured using mist nets. Birds with a brood patch were released immediately. In total, 42 adult waxbills were withheld for respirometry analyses. Upon capture, these birds were tagged with colored plastic rings for individual identification, aged, sexed, and then brought to the Animal House of the University of KwaZulu-Natal, Pietermaritzburg, where they were housed in outdoor aviaries (1 x 3 x 2 m) with shelter from sun and rain and with perches provided. Food (finch mix and millet) and water were provided ad libitum. Measurements of metabolic rate in the nocturnal ρ-phase started on the same day as the birds were captured. Birds were kept in the aviary for an average of 10 days and released at the original capture site. This work was carried out under ethical permit ‘VIB EC2022-090’, SAFRING permit No. ‘0163’ and Ezemvelo KZN Wildlife permit ‘OP 475’.

### ρ-(resting) phase metabolic rate

At sunset (19h00), after being weighed to the nearest 0.1 g, birds were placed in a 1.1 L airtight plastic chamber within a temperature-controlled environmental chamber (CMP2244, Conviron, Winnipeg, Canada) for ρ-phase metabolic rate measurement. The six temperatures tested were 12, 16, 20, 24, 28, and 31.5°C. Each plastic chamber contained a chicken-wire mesh to ensure a normal sleeping posture. Metabolic rate was estimated using flow-through respirometry (Lighton, 2018) by measuring O_2_ consumption (VO_2_; ml/min). Up to seven birds were measured during the same night. Ambient air was supplied by two pumps and divided into separate streams directed to a mass-flow meter (FB-8, Sable Systems) to provide a constant flow of ∼650 ml/min. Excurrent air from the bird and the baseline channels was alternately subsampled and passed through a Field Metabolic System (FMS-3, Sable Systems). Birds were measured alternately in cycles along with multiple baselines. The time of measurement for each bird within a cycle (and the length of each cycle) depended on the number of birds within a session. The average measurement time was 11 h. The first 2 h were discarded to ensure that birds were post-absorptive. After the respirometry measurement (06h00), the birds were weighed again to the nearest 0.1 g and returned to the aviary. Some individuals were measured more than once at certain temperatures, resulting in a dataset with repeated measurements. Further details of the respirometry setup can be found in Pacioni et al. (2023).

### α-(active) phase metabolic rate

To determine α-phase metabolic rates, birds were first weighed to the nearest 0.1 g, and then placed in a 1.1 L airtight plastic chamber within the temperature-controlled environmental chamber (see above). The temperature in this chamber was maintained at 28°C (which is within the TNZ of the South African waxbill, see below). Each day, four birds were measured using the same respirometry set up described above, in 5-min cycles, for a total of 10 cycles per bird.

The birds were kept in the metabolic chambers from 14h00 to 18h00. After the respirometry measurements, the birds were weighed again to the nearest 0.1 g and returned to the aviary. The same four birds were not selected for the ρ-phase metabolic rates on that night to reduce potential stress.

### Respirometry and data analyses

We used ExpeData software (Sable Systems) to record each experimental trial and extract metabolic rate values (ml O_2_/min). To estimate both the ρ-phase and the α-phase metabolic rates, the lowest stable section of the curve (averaged over 5 min) was selected using equation 9.7 from Lighton (2018).

The effect of ambient temperature (12, 16, 20, 24, 28, and 31.5°C) on ρ-phase metabolic rates was analyzed using a generalized additive mixed model (GAMM), with individual bird ID specified as a random effect (’gamm’ function from ‘mgcv’ R-package; Wood, 2023). A segmented regression model was applied to identify points at which the relationship between ρ-phase metabolic rates and temperature may change (’segmented’ R-package; Muggeo, 2008). Changes in α-phase metabolic rates (28°C) were tested by quantifying the variation in metabolic rate over time using measurements taken at 5-min intervals over 4 h. A GAMM was used to identify temporal trends in α-phase metabolic rates at 28°C. The inflection point of the GAMM curve was identified to delineate the period distinguishing ‘fasted’ and ‘non-fasted’ stage. Consequently, linear regression models (‘lm’ function) with a Gaussian error distribution were used to investigate whether the metabolic rates during the ρ-phase (28°C) differed significantly from those during the fasted α-phase (28°C). When repeated measurements per individual were available, the lowest metabolic rate per individual was selected. The GAMM (’gamm’ function) incorporated a basis dimension (k) of 3 as a value for the smooth terms in the GAMM fits to capture relationships no more complex than unimodal, and included individual ID as a random effect (Wood, 2023).

Sex was included as an independent factor in all models. Models were first run using whole-body metabolic rates and then using mass-independent metabolic rates, determined as the residuals derived from regressions of (log-transformed) ρ-phase and (log-transformed) α-phase metabolic rates on (log-transformed) body mass. Three different measures of body mass were considered in the analysis: body mass before and after the metabolic rate measurements, and the mean of these two measures. As consistent results were obtained for all three measures, the body mass measured after the metabolic measurements was used in all subsequent analyses. Variables and their interaction that were not statistically significant were removed using a backward stepwise procedure. Given that birds were caged for several days, number of days since capture was included as a fixed covariate to account for the potential effect of captivity. Interquartile ranges were used as a criterion to identify outliers (14 outliers were identified in the α-phase metabolic rates dataset), using the ‘quantile’ function in R. Shapiro-Wilk tests were performed to check the normality of the residuals (Shapiro-Wilk W > 0.9). The significance level was set at p ≤ 0.05. Body mass, ρ-phase, and α-phase metabolic rates were log transformed before analyses. All statistical analyses were performed using R software v. 4.2.2 (R Core Team 2022), details of which are available in the Supplementary File (RMarkdown file).

## Results

### ρ-phase metabolic rates

The values for ρ-phase metabolic rates are summarized in Table 1. Birds exhibited their lowest whole-body and mass-independent rates when exposed to an ambient temperature of 28°C, and showed an increase in metabolic rate below this temperature (whole-body, Figure 1A; mass-independent, Supplementary information Figure S1A). As the temperature decreased to ∼16°C (inflection point), metabolic rates also began to decrease (Figure 1B and Supplementary information Figure S1B).

**Figure 1.**
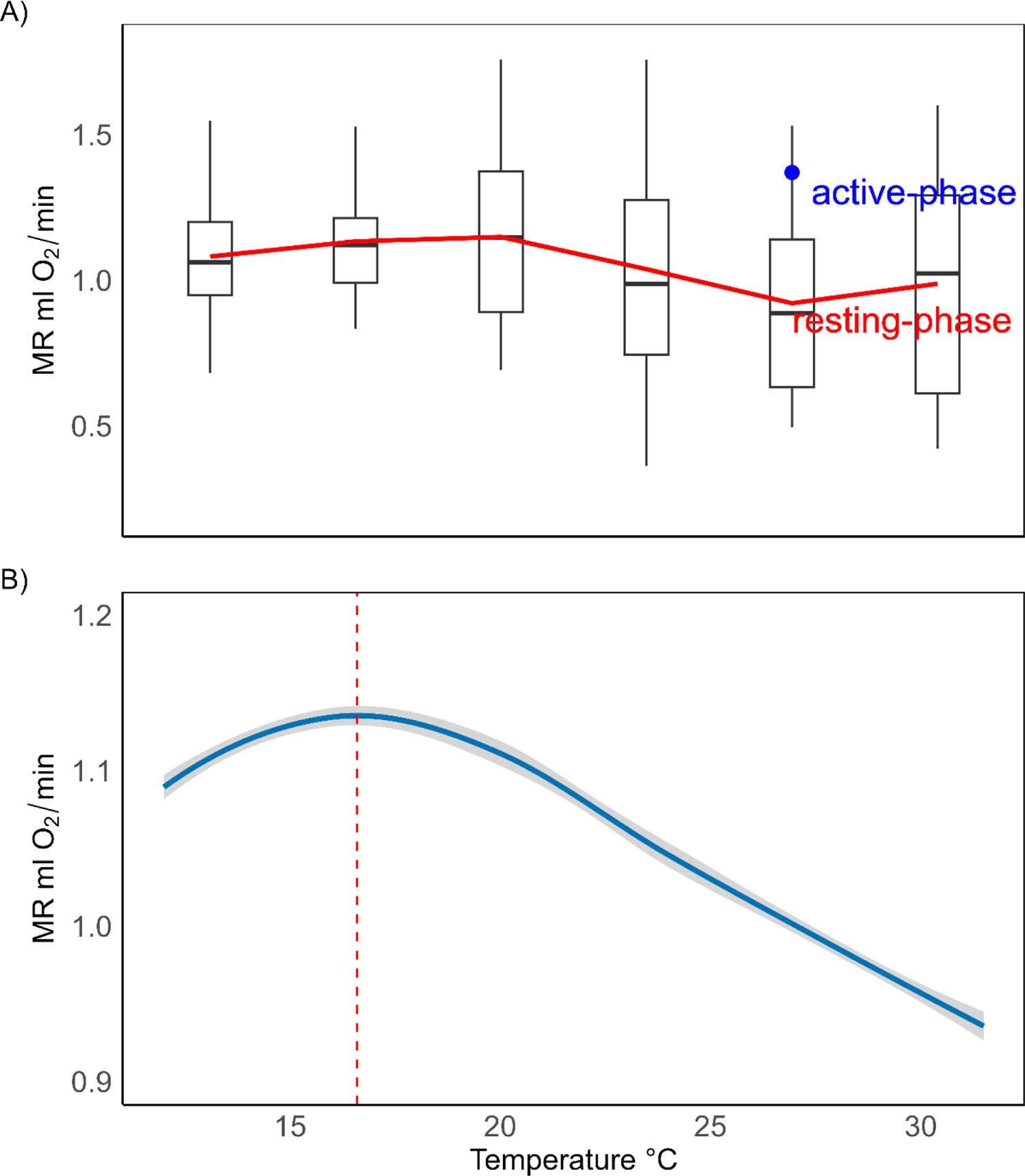
A) Boxplots of whole-body ρ-(resting) phase metabolic rate (ml O_2_/min) of wild common waxbill (*Estrilda astrild*) at various ambient temperatures. (Note: The blue filled circle represents the mean value of the α-(active) phase metabolic rates measured at 28°C. Boxplot whiskers extend to the minimum or maximum value within 1.5 times the interquartile range). B) The whole-body ρ-(resting) phase metabolic rate (MR ml O_2_/min) at various temperatures. (Note: The vertical red dashed line indicates the inflection point at 16.6°C. The shaded band indicates the 95% confidence interval around the segmented regression line, highlighting the region where there is 95% confidence in the true regression line).

**Table 1.** Mean ± SD (standard deviation), minimum-maximum values, and sample size (*n*) of the ρ-(resting) phase whole-body metabolic rates (ml O_2_/min) for various temperatures.

| **ρ- (resting) phase metabolic rate** |  |  |  |
| --- | --- | --- | --- |
| **Temperature (°C)** | **Mean ± SD**<br>**(ml O<sub>2</sub>/min)** | **Min-Max**<br>**(ml O<sub>2</sub>/min)** | **$n$** |
| **12** | 1.08 ± 0.21 | 0.68 - 1.55 | 27 |
| **16** | 1.13 ± 0.19 | 0.83 - 1.63 | 27 |
| **20** | 1.15 ± 0.28 | 0.69 - 1.76 | 29 |
| **24** | 1.04 ± 0.36 | 0.36 - 1.76 | 28 |
| **28** | 0.92 ± 0.31 | 0.50 - 1.53 | 49 |
| **31.5** | 0.99 ± 0.40 | 0.42 - 1.60 | 19 |

### α-phase metabolic rates

GAMM analyses supported a decline in metabolic rate over time, for both whole-body (p < 0.001; Figure 2) and mass-independent (p < 0.001; Figure S2) measurements, visually plateauing off at ∼120 min (Figure 2 and Supplementary information Figure S2). Whole-body α-phase metabolic rates (fasted birds only) were on average 48% higher than whole-body ρ-phase metabolic rates (28°C), and this difference was also significant for both whole-body (p > 0.05, Figure 1A) and mass-independent (p > 0.05, Supplementary information Figure S1A) measurements. There was no evidence of a positive correlation between ρ-phase metabolic rate (28°C) and α-phase metabolic rate (fasted birds only). Neither sex nor number of days since capture were significant (all p > 0.10).

**Figure 2.**
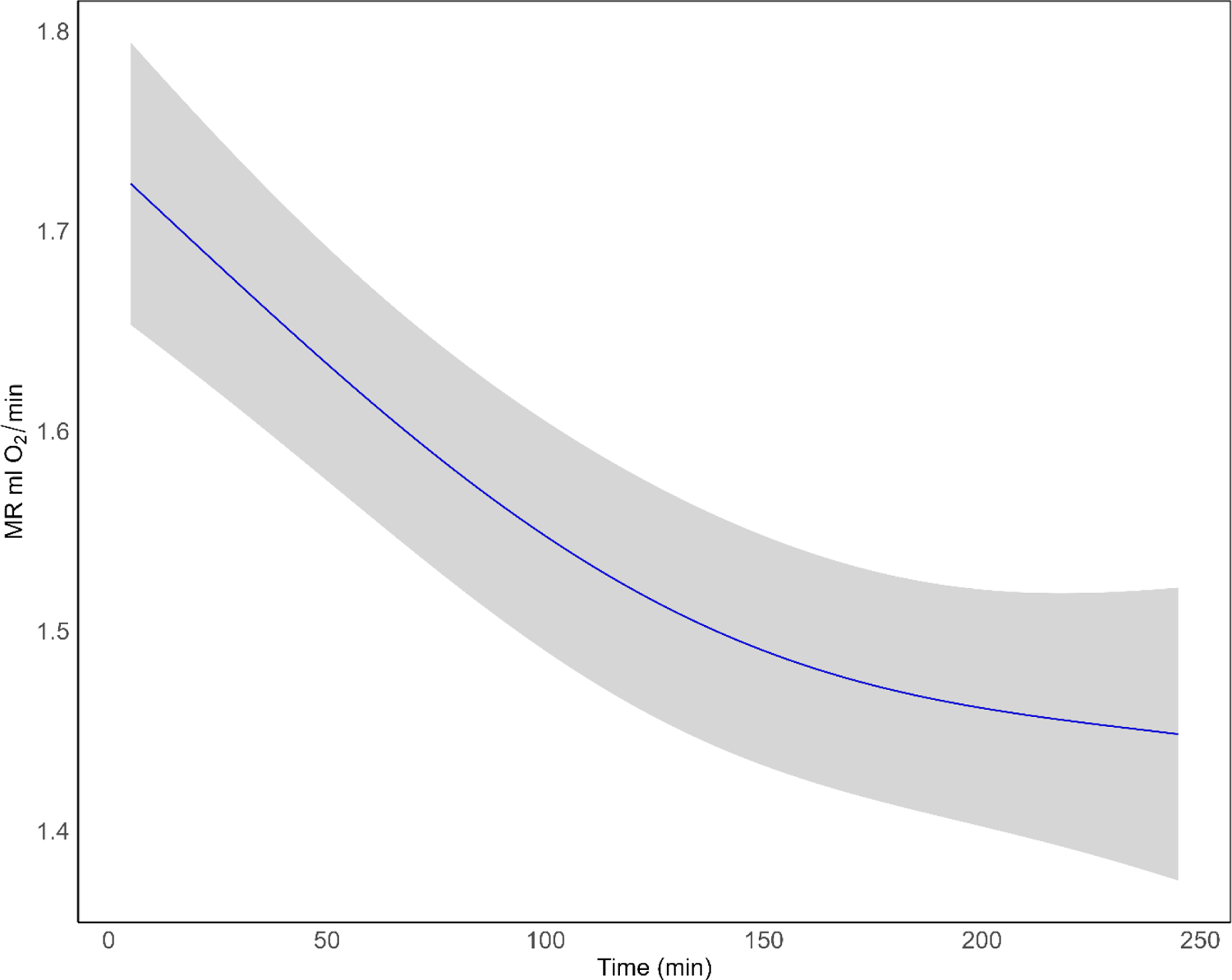
Whole-body α-(active) phase metabolic rates (ml O_2_/min) of common waxbill (*Estrilda astrild*) over the 4 h fasting period. (Note: The shaded ribbon represents the 95% confidence interval).

## Discussion

The observed thermoneutral zone (TNZ) curve predominantly follows the classic Scholander-Irving model, where endotherms maintain a stable basal metabolic rate (BMR) within a range of ambient temperatures, but must produce additional metabolic energy to keep their body temperature constant when the ambient temperature falls below or rises above it. Common waxbill metabolic rates were lowest at an ambient temperature of 28°C. Below this temperature, ρ-phase (resting) metabolic rates increased linearly, until an inflection point at around 16°C where birds began to decrease their ρ-phase metabolic rates. We also found that α-(active) phase metabolic rates (fasted) were 48% higher than ρ-phase metabolic rates (28°C), and that whole-body and mass-independent α-phase metabolic rates showed an unimodal decrease over time, plateauing off at ∼120 min.

Although the specific limits of the TNZ could not be precisely defined because of a temperature difference of 4°C between the ambient temperatures tested, it is reasonable to conclude that the lower value of the TNZ (i.e., the lower critical temperature) for common waxbills fell approximately around 28°C. This was consistent with findings from captive individuals of this species (Pacioni et al., 2023). As temperatures fell below this point, we observed an increase in both whole-body and mass-independent ρ-phase metabolic rates. However, the curve showed an inflection point at around 16°C, where both whole-body and mass-independent ρ-phase metabolic rates began to decrease. This may suggest adaptive mechanisms, such as facultative hypothermic responses, to conserve energy in colder conditions (Geiser, 2021; McKechnie & Lovegrove, 2002). Facultative hypothermia, such as resting phase hypothermia, refers to the adaptive ability to reduce body temperature in response to environmental conditions (e.g., cold temperatures), with a 30-40% reduction in metabolic rate (Reinertsen, 1983). This phenomenon has been observed in species spanning 16 different orders and 31 families of birds (Ritchison, 2023), some of which have a small body mass (Ruf & Geiser, 2015). For example, small passerines (i.e., blue tit *Cyanistes caeruleus*, willow tit *Poecile montanus*, and black-capped chickadee *Poecile atricapillus*) have demonstrated the ability to reduce their body temperature by several degrees after prolonged exposure to cold conditions as an energy-saving strategy to increase winter survival (Brodin et al., 2017). However, a study by Andreasson et al. (2019) found that juvenile great tits (*Parus major*) and juvenile blue tits were less likely to adopt hypothermia as an energy-saving strategy at very low ambient temperatures (∼-8°C) when faced with an increased perceived predation risk. Similarly, Cooper and Gessaman (2005) showed that mountain chickadees (*Poecile gambeli*) and juniper titmice (*Baeolophus ridgwayi*) used more energy to reduce their body temperature when the ambient temperature was exceptionally low (∼-10°C). As the temperatures examined in our study were not extremely low, a potential hypothermic mechanism is likely to be energetically advantageous for our birds.

Numerous studies have shown a breakdown of the Scholander-Irving model at lower temperatures, with metabolic rates deviating from linearity at lower temperatures (Noakes et al., 2013; Nzama et al., 2010; Steiger et al., 2009; Thabethe et al., 2013; van de Ven et al., 2013). However, these studies did not discuss the precise mechanism underlying this metabolic downregulation at lower temperatures. As we did not monitor body temperature during our measurements, it is important to acknowledge this limitation of our study. Without direct measurements of body temperature, our understanding of the observed inflection point in our TNZ curve is limited, leading to speculation and highlighting a potential avenue for future research into the relationship between body temperature adaptations and responses to colder temperatures in our study species.

As the physiological ability to adapt to low temperatures has been shown to influence the geographic distribution of endothermic organisms (Khaliq et al., 2017; Hayes et al., 2018), range distribution expansion may be limited by physiological constraints, such as those associated with the ability to sustain energetically elevated rates of thermogenesis (heat production) for long periods (Buckley et al., 2018). Indeed, although cold temperatures outside the TNZ are not directly lethal to the bird itself, the prolonged maintenance of high metabolic rates, and hence high energy requirements, may be constrained by environmental factors (e.g., food availability). This is particularly relevant for invasive species such as the common waxbill, which tends to occupy colder climates in its invasive distribution range compared with its native distribution range (Stiels et al., 2011). Lasiewski et al. (1964) and Stephens et al. (2001) conducted comparable studies on black-rumped waxbill and orange-cheeked waxbill, both of which have invasive populations in Iberia along with the common waxbill. These studies examined the TNZ of these species at ambient temperatures similar to or lower than those in our study. Their results showed a linear up-regulation of metabolic rate when the ambient temperatures were below the TNZ for both species. This evidence suggests that these two species may lack the ability to limit energy expenditure at colder temperatures and may explain why the common waxbill is a more successful invasive species in Iberia than its counterparts (Ascensão et al., 2021).

In addition to TNZ, we also examined the patterns of α-phase (active) metabolic rates and found significant differences between the (whole-body and mass-independent) ρ-phase (28°C) and the α-phase (28°C, fasted birds only) metabolic rates. Specifically, during the α-phase, the whole-body metabolic rates of the fasted birds were significantly higher by 48% than during the ρ-phase. These results do not support the conclusion of Lasiewski et al. (1964) and Stephens et al. (2001) that metabolic rates during ρ-phase and α-phase are generally not different in small birds. Furthermore, a recent study by Ellis and Gabrielsen (2019) argued that both α-phase and ρ-phase metabolic rates represent BMR. Other studies that have found evidence of a circadian rhythm in metabolic rates have typically reported smaller differences. For example, McKechnie and Lovegrove (1999) found that the ρ-phase BMR was approximately 20% lower than the α-phase BMR in fed black-shouldered kites (*Elanus caeruleus*). Daan et al. (1989) found that, in kestrels (*Falco tinnunculus*), the metabolic rate during the α-phase was 22-27% higher than the metabolic rate during the ρ-phase. However, these studies involved metabolic measurements lasting for 24 h. The longer measurement period may have resulted in a lower percentage increase in metabolic rate during the α-phase compared with the 4 h measurements we performed. This difference could be attributed to potential acclimatization effects, where birds might adjust to the laboratory conditions over time, leading to a lower metabolic response during the α-phase. The duration of exposure may then influence the degree to which birds modulate their metabolic rates, highlighting the importance of considering the duration of measurements when understanding circadian rhythm-related variations in avian energetics.

Finally, we observed an effect of fasting on the α-phase metabolic rates, with a statistically significant decrease in metabolic rates from a non-fasted to a fasted state. This reduction in metabolism is likely to be a consequence of the energy expenditure associated with digesting and assimilating an ingested meal (Brody & Lardy, 1946; Rubner, 1902). Similarly, Cade et al. (1965) observed a decrease in CO_2_ production in the black-rumped waxbill during the transition from fed to fasting conditions, with α-phase metabolic rate values at 3 h post-feeding approximately 30% lower than initial values. Furthermore, in line with these results, we found that common waxbills reached a post-absorptive state at ∼120 min, as shown in Figure 2, where metabolic rate consumption started to plateau, with final metabolic rate values about 40% lower than the initial values. These values suggest a rapid digestion process, consistent with other estrildid species (Cade et al., 1965).

## Conclusions

Our results highlight the potential use of an energy-saving strategy by common waxbills as an adaptive response to cold, which may explain their success as an invasive species. Although metabolic inflection points have been repeatedly identified in studies of TNZ, the specific mechanisms underlying this metabolic down-regulation at low temperatures have not been investigated by many authors. Therefore, we suggest that future research should prioritize the study of body temperature variation, focusing on elucidating its potential contribution to metabolic adaptation to colder environments. Finally, we found that metabolic rates in the α-phase were significantly higher than those in the ρ-phase. Therefore, we do not support the conclusion of Lasiewski et al. (1964) and Stephens et al. (2001) that metabolic rates during α-phase and ρ-phase are generally not different in small birds.

## Funding and Acknowledgements

We thank Preshnee Singh and Ebrahim Ally (Centre for Functional Biodiversity, School of Life Sciences, University of KwaZulu-Natal) for their invaluable assistance during the fieldwork. This study was supported by the Research Foundation - Flanders (grant #G0E4320 N) through a bilateral research collaboration with the Russian Science Foundation (grant #20-44-01005). In addition, the Methusalem Project 01M00221 (Ghent University), awarded to Frederick Verbruggen, Luc Lens and An Martel, contributed to this research. Marina Sentís acknowledges the support of FWO-Vlaanderen (project 11E1623N). In addition, Cesare Pacioni and Marina Sentís acknowledge the support of two FWO travel grants (V441722N and V441822N), which made this study possible. Colleen T. Downs was supported by the National Research Foundation (ZA, grant 98404) and the University of KwaZulu-Natal.

## Declaration of competing interest

There were no conflicts of interest.

## Notes

https://doi.org/10.6084/m9.figshare.25020026.v1

### Competing Interest Statement

The authors have declared no competing interest.

## References

Allen, L. R., & Hume, I. D. (2001). The maintenance nitrogen requirement of the zebra finch *Taeniopygia guttata*. Physiological and Biochemical Zoology, 74(3), 366–375. 10.1086/320431

Andreasson, F., Nord, A., & Nilsson, J.-Å. (2019). Age-dependent effects of predation risk on night-time hypothermia in two wintering passerine species. Oecologia, 189(2), 329–337. 10.1007/s00442-018-04331-7

Ascensão, F., D’Amico, M., Martins, R. C., Rebelo, R., Barbosa, A. M., Bencatel, J., … & Capinha, C. (2021). Distribution of alien tetrapods in the Iberian Peninsula. NeoBiota, 64, 1–21. 10.3897/neobiota.64.55597

Aschoff, J., & Pohl, H. (1970). Rhythmic variations in energy metabolism. Federation Proceedings, 29(4), 1541–1552.

Beaulieu, M. (2016). A bird in the house: the challenge of being ecologically relevant in captivity. Frontiers in Ecology and Evolution, 4, 141. https://www.frontiersin.org/articles/10.3389/fevo.2016.00141

Boardman, L., Lockwood, J. L., Angilletta Jr, M. J., Krause, J. S., Lau, J. A., Loik, M. E., … & Meyerson, L. A. (2022). The future of invasion science needs physiology. BioScience, 72(12), 1204–1219. 10.1093/biosci/biac080

Bongaarts, J. (2019). IPBES, 2019. Summary for policymakers of the global assessment report on biodiversity and ecosystem services of the Intergovernmental Science-Policy Platform on Biodiversity and Ecosystem Services. Population and Development Review, 45(3), 680–681. 10.1111/padr.12283

Bozinovic, F., Calosi, P., & Spicer, J. I. (2011). Physiological Correlates of Geographic Range in Animals. Annual Review of Ecology, Evolution, and Systematics, 42(1), 155–179. 10.1146/annurev-ecolsys-102710-145055

Brodin, A., Nilsson, J.-Å., & Nord, A. (2017). Adaptive temperature regulation in the little bird in winter: Predictions from a stochastic dynamic programming model. Oecologia, 185(1), 43–54. 10.1007/s00442-017-3923-3

Brody, S., & Lardy, H. A. (1946). Bioenergetics and Growth. The Journal of Physical Chemistry, 50(2), 168–169. 10.1021/j150446a008

Buckley, L. B., Khaliq, I., Swanson, D. L., & Hof, C. (2018). Does metabolism constrain bird and mammal ranges and predict shifts in response to climate change? Ecology and Evolution, 8(24), 12375–12385. 10.1002/ece3.4537

Cade, T. J., Tobin, C. A., & Gold, A. (1965). Water economy and metabolism of two estrildine finches. Physiological Zoology, 38(1), 9–33. 10.1086/physzool.38.1.30152342

Cardoso, G. C., & Reino, L. (2018). Ecologically benign invasions: The invasion and adaptation of common waxbills (*Estrilda astrild*) in Iberia. Histories of Bioinvasions in the Mediterranean, 149–169. 10.1007/978-3-319-74986-0_7

Castro-Díez, P., Vaz, A. S., Silva, J. S., Van Loo, M., Alonso, Á., Aponte, C., Bayón, Á., Bellingham, P. J., Chiuffo, M. C., DiManno, N., Julian, K., Kandert, S., La Porta, N., Marchante, H., Maule, H. G., Mayfield, M. M., Metcalfe, D., Monteverdi, M. C., Núñez, M. A., … Godoy, O. (2019). Global effects of non-native tree species on multiple ecosystem services. Biological Reviews, 94(4), 1477–1501. 10.1111/brv.12511

Cooper, C. E., Hurley, L. L., Deviche, P., & Griffith, S. C. (2020). Physiological responses of wild zebra finches (*Taeniopygia guttata*) to heatwaves. Journal of Experimental Biology, 223(12), jeb225524. 10.1242/jeb.225524

Cooper, C. E., Withers, P. C., Hurley, L. L., & Griffith, S. C. (2019). The field metabolic rate, water turnover, and feeding and drinking behavior of a small avian desert granivore during a summer heatwave. Frontiers in Physiology, 1405. https://www.frontiersin.org/articles/10.3389/fphys.2019.01405

Cooper, S. J., & Gessaman, J. A. (2005). Nocturnal hypothermia in seasonally acclimatized mountain chickadees and juniper titmice. The Condor, 107(1), 151–155. 10.1093/condor/107.1.151

Daan, S., Masman, D., Strijkstra, A., & Verhulst, S. (1989). Intraspecific allometry of basal metabolic rate: relations with body size, temperature, composition, and circadian phase in the kestrel, *Falco tinnunculus*. Journal of Biological Rhythms, 4(2), 155–171. 10.1177/074873048900400212

Diagne, C., Leroy, B., Vaissière, A.-C., Gozlan, R. E., Roiz, D., Jarić, I., Salles, J.-M., Bradshaw, C. J. A., & Courchamp, F. (2021). High and rising economic costs of biological invasions worldwide. Nature, 592(7855), 571–576. 10.1038/s41586-021-03405-6

Dukes, J. S., & Mooney, H. A. (1999). Does global change increase the success of biological invaders? Trends in Ecology & Evolution, 14(4), 135–139. 10.1016/S0169-5347(98)01554-7

Dyer, E. E., Redding, D. W., & Blackburn, T. M. (2017). The global avian invasions atlas, a database of alien bird distributions worldwide. Scientific Data, 4(1), Article 1. 10.1038/sdata.2017.41

Ellis, H. I., & Gabrielsen, G. W. (2019). Reassessing the definition of basal metabolic rate: Circadian considerations in avian studies. Comparative Biochemistry and Physiology Part A: Molecular & Integrative Physiology, 237, 110541. 10.1016/j.cbpa.2019.110541

Geiser, F. (2021). Ecological Physiology of Daily Torpor and Hibernation. Springer International Publishing. 10.1007/978-3-030-75525-6

Gerson, A. R., Cristol, D. A., & Seewagen, C. L. (2019). Environmentally relevant methylmercury exposure reduces the metabolic scope of a model songbird. Environmental Pollution, 246, 790–796. 10.1016/j.envpol.2018.12.072

Gleditsch, J. M., & Brooks, D. M. (2020). Scaly-breasted Munia (*Lonchura punctulata* Linnaeus 1758). Invasive Birds: Global Trends and Impacts, 159–162. 10.1079/9781789242065.0159

Goodwin, D. (1982). Estrildid Finches of the World (First Edition). British Museum, London.

Hayes, J. P., Feldman, C. R., & Araújo, M. B. (2018). Mass-independent maximal metabolic rate predicts geographic range size of placental mammals. Functional Ecology, 32(5), 1194–1202. 10.1111/1365-2435.13053

Hellmann, J. J., Byers, J. E., Bierwagen, B. G., & Dukes, J. S. (2008). Five potential consequences of climate change for invasive species. Conservation biology, 22(3), 534–543. 10.1111/j.1523-1739.2008.00951.x

Huey, R. B., Ma, L., Levy, O., & Kearney, M. R. (2021). Three questions about the eco-physiology of overwintering underground. Ecology Letters, 24(2), 170–185. 10.1111/ele.13636

Khaliq, I., Böhning-Gaese, K., Prinzinger, R., Pfenninger, M., & Hof, C. (2017). The influence of thermal tolerances on geographical ranges of endotherms. Global Ecology and Biogeography, 26(6), 650–668. 10.1111/geb.12575

Khaliq, I., Hof, C., Prinzinger, R., Böhning-Gaese, K., & Pfenninger, M. (2014). Global variation in thermal tolerances and vulnerability of endotherms to climate change. Proceedings of the Royal Society B: Biological Sciences, 281(1789), 20141097. 10.1098/rspb.2014.1097

Lasiewski, R. C., Hubbard, S. H., & Moberly, W. R. (1964). Energetic relationships of a very small passerine bird. The Condor, 66(3), 212–220. 10.2307/1365646

Lever C. (2005). Naturalised Birds of the World. A&C Black. https://www.nhbs.com/naturalised-birds-of-the-world-book

Lighton, J. R. B. (2018). Measuring Metabolic Rates: A Manual for Scientists (2nd ed.). Oxford University Press. 10.1093/oso/9780198830399.001.0001

Liu, C., Wolter, C., Courchamp, F., Roura-Pascual, N., & Jeschke, J. M. (2022). Biological invasions reveal how niche change affects the transferability of species distribution models. Ecology, 103(8), e3719. 10.1002/ecy.3719

Mainka, S. A., & Howard, G. W. (2010). Climate change and invasive species: Double jeopardy. Integrative Zoology, 5(2), 102–111. 10.1111/j.1749-4877.2010.00193.x

Marschall, U., & Prinzinger, R. (1991). Vergleichende Ökophysiologie von fünf Prachtfinkenarten (Estrildidae). Journal für Ornithologie, 132(3), 319–323. 10.1007/BF01640540

McKechnie, A. E., & Lovegrove, B. G. (1999). Circadian metabolic responses to food deprivation in the black-shouldered kite. The Condor, 101(2), 426–432. 10.2307/1370010

McKechnie, A. E., & Lovegrove, B. G. (2002). Avian facultative hypothermic responses: a review. The Condor, 104(4), 705–724. 10.1093/condor/104.4.705

McNab, B. K. (2012). Extreme Measures: The Ecological Energetics of Birds and Mammals. University of Chicago Press.

Milne, R., Cunningham, S. J., Lee, A. T. K., & Smit, B. (2015). The role of thermal physiology in recent declines of birds in a biodiversity hotspot. Conservation Physiology, 3(1), cov048. 10.1093/conphys/cov048

Muggeo, V. M. (2008). Segmented: an R package to fit regression models with broken-line relationships. R news, 8(1), 20–25.

Noakes, M. J., Smit, B., Wolf, B. O., & McKechnie, A. E. (2013). Thermoregulation in African Green Pigeons (*Treron calvus*) and a re-analysis of insular effects on basal metabolic rate and heterothermy in columbid birds. Journal of Comparative Physiology B, 183(7), 969–982. 10.1007/s00360-013-0763-2

Nzama, S. N., Downs, C. T., & Brown, M. (2010). Seasonal variation in the metabolism-temperature relation of House Sparrows (*Passer domesticus*) in KwaZulu-Natal, South Africa. Journal of Thermal Biology, 35(2), 100–104. 10.1016/j.jtherbio.2009.12.002

Pacioni, C., Sentís, M., Kerimov, A., Bushuev, A., Lens, L., & Strubbe, D. (2023). Seasonal variation in thermoregulatory capacity of three closely related Afrotropical Estrildid finches introduced to Europe. Journal of Thermal Biology, 113, 103534. 10.1016/j.jtherbio.2023.103534

Pyšek, P., Hulme, P. E., Simberloff, D., Bacher, S., Blackburn, T. M., Carlton, J. T., … & Richardson, D. M. (2020). Scientists’ warning on invasive alien species. Biological Reviews, 95(6), 1511–1534. 10.1111/brv.12627

Reinertsen, R. E. (1983). Nocturnal hypothermia and its energetic significance for small birds living in the arctic and subarctic regions. A review. Polar Research, 1(3), 269–284. 10.1111/j.1751-8369.1983.tb00743.x

Reino, L., Figueira, R., Beja, P., Araújo, M. B., Capinha, C., & Strubbe, D. (2017). Networks of global bird invasion altered by regional trade ban. Science Advances, 3(11), e1700783. 10.1126/sciadv.1700783

Ribeiro, J., Sillero, N., Lopes, R. J., Sullivan, M. J. P., Santana, J., Capinha, C., & Reino, L. (2020). Common Waxbill (Estrilda astrild Linnaeus, 1758). Invasive Birds: Global Trends and Impacts, 155–158. 10.1079/9781789242065.0155

Ritchison, G. (2023). Energy Balance and Thermoregulation. In G. Ritchison (Ed.), In a Class of Their Own: A Detailed Examination of Avian Forms and Functions (pp. 1253–1401). Springer International Publishing. 10.1007/978-3-031-14852-1_10

Ros, M., Lacerda, M. B., Vázquez-Luis, M., Masunari, S., & Guerra-García, J. M. (2016). Studying exotics in their native range: Can introduced fouling amphipods expand beyond artificial habitats?. Biological Invasions, 18, 2983–3000. 10.1007/s10530-016-1191-5

Rubner, M. (1902). Die Gesetze des Energieverbrauchs bei der Ernährung. Franz Deuticke. Leipzig & Vienna.

Ruf, T., & Geiser, F. (2015). Daily torpor and hibernation in birds and mammals. Biological Reviews, 90(3), 891–926. 10.1111/brv.12137

Scholander, P. F., Hock, R., Walters, V., & Irving, L. (1950). Adaptation to cold in arctic and tropical mammals and birds in relation to body temperature, insulation, and basal metabolic rate. The Biological Bulletin, 99(2), 259–271. 10.2307/1538742

Sentís, M., Pacioni, C., De Cuyper, A., Janssens, G. P. J., Lens, L., & Strubbe, D. (2023). Biophysical models accurately characterize the thermal energetics of a small invasive passerine bird. iScience, 26(10), 107743. 10.1016/j.isci.2023.107743

Sheldon, E. L., & Griffith, S. C. (2018). Embryonic heart rate predicts prenatal development rate, but is not related to post-natal growth rate or activity level in the zebra finch (*Taeniopygia guttata*). Ethology, 124(11), 829–837. 10.1111/eth.12817

Shirley, S. M., & Kark, S. (2009). The role of species traits and taxonomic patterns in alien bird impacts. Global Ecology and Biogeography, 18(4), 450–459. 10.1111/j.1466-8238.2009.00452.x

Steiger, S. S., Kelley, J. P., Cochran, W. W., & Wikelski, M. (2009). Low metabolism and inactive lifestyle of a tropical rain forest bird investigated via heart-rate telemetry. Physiological and Biochemical Zoology, 82(5), 580–589. 10.1086/605336

Stephens, C. M., Siegel, R. B., & Weathers, W. W. (2001). Thermal conductance and basal metabolism of the Orange-cheeked Waxbill (*Estrilda melpoda*). Ostrich, 72(1–2), 121–123. 10.2989/00306520109485299

Stiels, D., Gaißer, B., Schidelko, K., Engler, J. O., & Rödder, D. (2015). Niche shift in four non-native estrildid finches and implications for species distribution models. Ibis, 157(1), 75–90. 10.1111/ibi.12194

Stiels, D., Schidelko, K., Engler, J. O., Van Den Elzen, R., & Rödder, D. (2011). Predicting the potential distribution of the invasive Common Waxbill *Estrilda astrild* (Passeriformes: Estrildidae). Journal of Ornithology, 152(3), 769–780. 10.1007/s10336-011-0662-9

Strubbe, D., Jackson, H., Groombridge, J., & Matthysen, E. (2015). Invasion success of a global avian invader is explained by within-taxon niche structure and association with humans in the native range. Diversity and Distributions, 21(6), 675–685. 10.1111/ddi.12325

Strubbe, D., Jiménez, L., Barbosa, A. M., Davis, A. J. S., Lens, L., & Rahbek, C. (2023). Mechanistic models project bird invasions with accuracy. Nature Communications, 14(1), 2520. 10.1038/s41467-023-38329-4

Stuart, K. C., Sherwin, W. B., Edwards, R. J., & Rollins, L. A. (2023). Evolutionary genomics: Insights from the invasive European starlings. Frontiers in Genetics, 13, 1010456. 10.3389/fgene.2022.1010456

Sullivan, M. J. P., & Franco, A. M. A. (2018). Changes in habitat associations during range expansion: Disentangling the effects of climate and residence time. Biological Invasions, 20(5), 1147–1159. 10.1007/s10530-017-1616-9

Thabethe, V., Thompson, L. J., Hart, L. A., Brown, M., & Downs, C. T. (2013). Seasonal effects on the thermoregulation of invasive rose-ringed parakeets (*Psittacula krameri*). Journal of Thermal Biology, 38(8), 553–559. 10.1016/j.jtherbio.2013.09.006

Urban, M. C. (2015). Accelerating extinction risk from climate change. Science, 348(6234), 571–573. 10.1126/science.aaa4984

van de Ven, T. M. F. N., Mzilikazi, N., & McKechnie, A. E. (2013). Seasonal Metabolic Variation in Two Populations of an Afrotropical Euplectid Bird. Physiological and Biochemical Zoology, 86(1), 19–26. 10.1086/667989

Walther, G.-R., Roques, A., Hulme, P. E., Sykes, M. T., Pyšek, P., Kühn, I., Zobel, M., Bacher, S., Botta-Dukát, Z., Bugmann, H., Czúcz, B., Dauber, J., Hickler, T., Jarošík, V., Kenis, M., Klotz, S., Minchin, D., Moora, M., Nentwig, W., … Settele, J. (2009). Alien species in a warmer world: Risks and opportunities. Trends in Ecology & Evolution, 24(12), 686–693. 10.1016/j.tree.2009.06.008

Winkler, D. W., Billerman, S. M., & Lovette, I. J. (2020). Waxbills and Allies (Estrildidae), version 1.0. Birds of the World. 10.2173/bow.estril1.01species_shared.bow.project_name

Wood, S. (2023). mgcv: Mixed GAM Computation Vehicle with Automatic Smoothness Estimation (1.9-0) [Computer software]. https://cran.r-project.org/web/packages/mgcv/index.html

