## Supplementary material for "Metabolic responses to cold: thermal physiology of native common waxbills (*Estrilda astrild*)"

### Supplementary material from "Metabolic responses to cold: thermal physiology of native common waxbills (*Estrilda astrild*)"

Cesare Pacioni[a*], Marina Sentís[a], Anvar Kerimov[b], Andrey Bushuev[b], Colleen T. Downs[c], Luc Lens[a], Diederik Strubbe[a]

[a] Terrestrial Ecology Unit, Ghent University, Ghent, Belgium

[b] Department of Vertebrate Zoology, Faculty of Biology, M.V. Lomonosov Moscow State University, Moscow, Russia

[c] Centre for Functional Biodiversity, School of Life Sciences, University of KwaZulu-Natal, Pietermaritzburg, South Africa

#### Abstract

Ecophysiological studies of invasive species tend to focus on captive individuals and their invasive range. However, the importance of gaining a thorough understanding of their physiology in their native range, where autecological knowledge is limited, has played a crucial role in assessing species ecophysiological responses to the often novel environmental conditions encountered in their invasive range. Here, we investigated the ecophysiological characteristics of a population of the wild-caught common waxbill (*Estrilda astrild*), a successful global invader, in part of its native range (South Africa). We investigated how this species adjusts its resting metabolic rate over a range of temperatures to identify its thermoneutral zone (TNZ) as an indicator of a species' thermal tolerance. The observed TNZ curve predominantly followed the classic Scholander-Irving model, with metabolic rates increasing linearly at temperatures outside the TNZ. However, we found an inflection point at moderately cold temperatures (16°C) where the common waxbill began to decrease its metabolic rate. This finding highlights the potential use of an energy-saving strategy as an adaptive response to cold, such as facultative hypothermic responses through a reduction in body temperature, which may explain their success as an invasive species. We argue that although metabolic infection points have been repeatedly identified in studies of TNZ, the specific mechanisms behind metabolic down-regulation at low temperatures remain underexplored in the literature. We therefore suggest that future research should focus on investigating body temperature variation, with particular emphasis on its potential contribution to metabolic adaptation in colder environments.

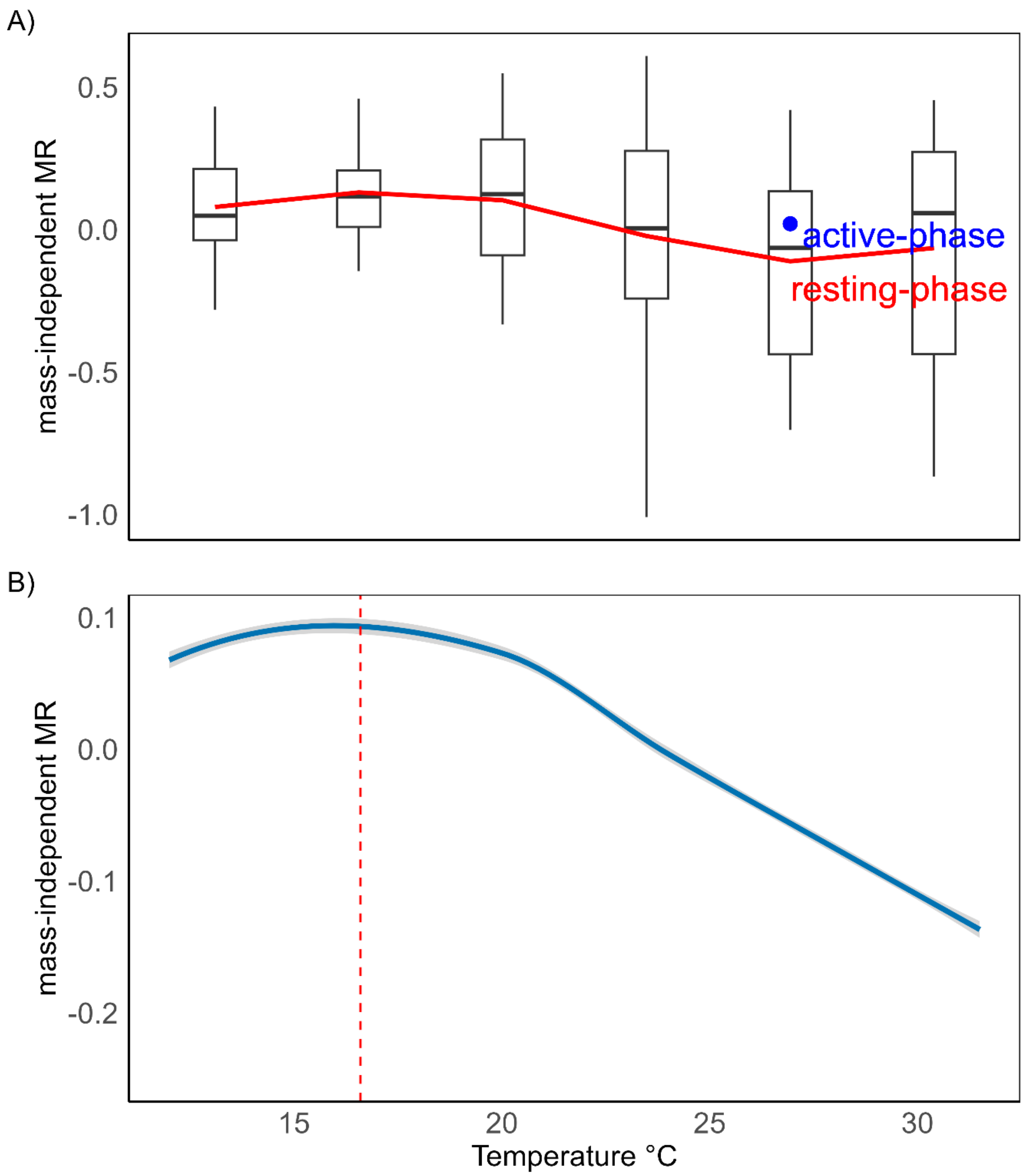


Figure S1. A) Boxplots of mass-independent ρ- (resting) phase metabolic rate of the common waxbill (*Estrilda astrild*) at various ambient temperatures. (Note: The blue filled circle represents the mean value of the α- (active) phase metabolic rates measured at 28°C. Boxplot whiskers extend to the minimum or maximum value within 1.5 times the interquartile range). B) The mass-independent ρ- (resting) phase metabolic rate at various temperatures. (Note: The vertical red dashed line indicates the inflection point at 16.6°C. The shaded band indicates the

95% confidence interval around the segmented regression line, highlighting the region where there is 95% confidence in the true regression line).

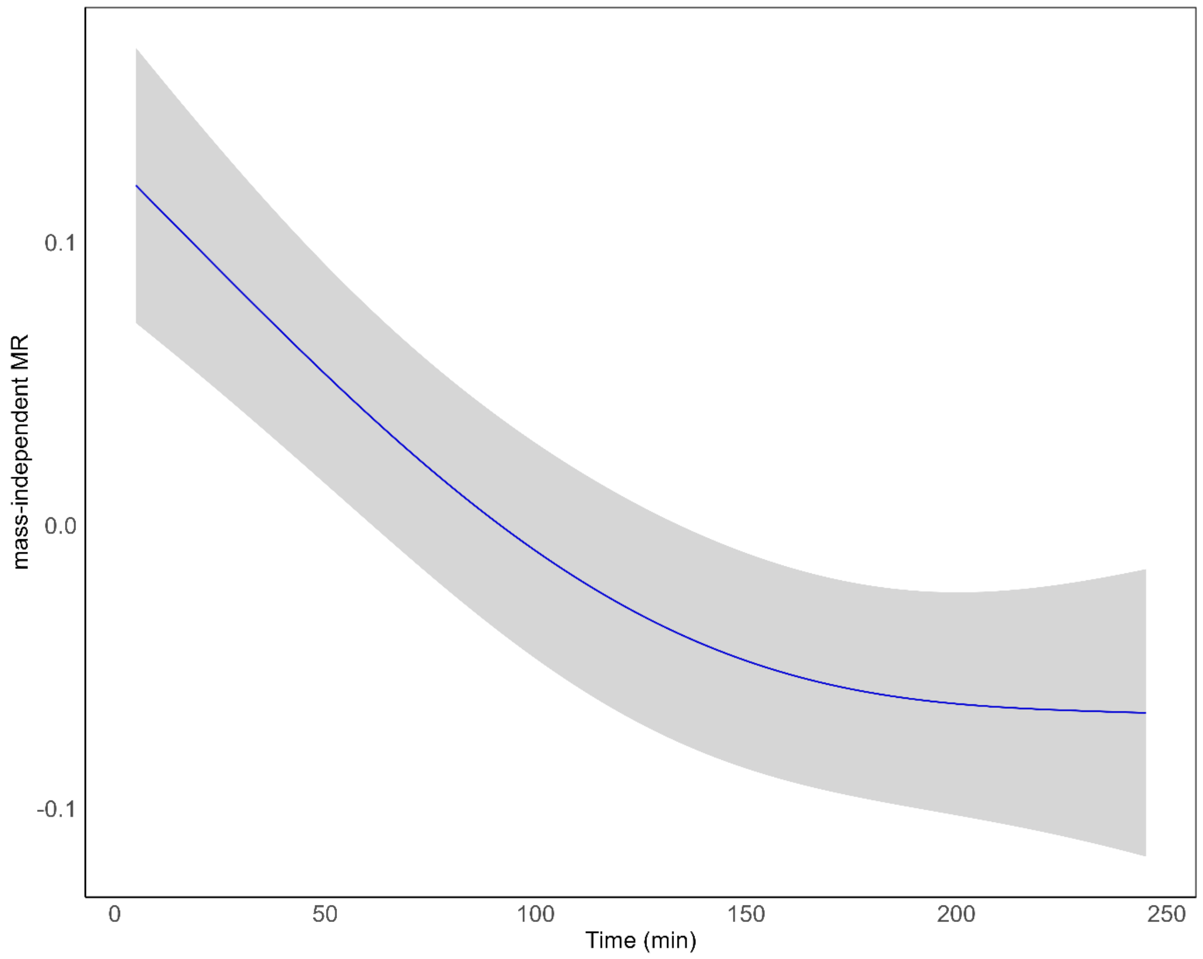

Figure S2. Mass-independent α- (active) phase metabolic rates of common waxbill (*Estrilda astrild*) over the four hour fasting period. (Note: The shaded ribbon represents the 95% confidence interval).
